# What is the test-retest reliability of common task-fMRI measures? New empirical evidence and a meta-analysis

**DOI:** 10.1101/681700

**Authors:** Maxwell L. Elliott, Annchen R. Knodt, David Ireland, Meriwether L. Morris, Richie Poulton, Sandhya Ramrakha, Maria L. Sison, Terrie E. Moffitt, Avshalom Caspi, Ahmad R. Hariri

**Author notes:** These authors contributed equally to this work (Maxwell L. Elliott, Annchen R. Knodt). Correspondence: Ahmad R. Hariri, Ph.D., Professor of Psychology and Neuroscience, Director, Laboratory of NeuroGenetics, Head, Cognition and Cognitive Neuroscience Training Program, Duke University, Durham, NC 27708, USA.

## Abstract

Identifying brain biomarkers of disease risk is a growing priority in neuroscience. The ability to identify meaningful biomarkers is limited by measurement reliability; unreliable measures are unsuitable for predicting clinical outcomes. Measuring brain activity using task-fMRI is a major focus of biomarker development; however, the reliability of task-fMRI has not been systematically evaluated. We present converging evidence demonstrating poor reliability of task-fMRI measures. First, a meta-analysis of 90 experiments (N=1,008) revealed poor overall reliability (mean ICC=.397). Second, the test-retest reliabilities of activity in *a priori* regions of interest across 11 common fMRI tasks collected in the context of the Human Connectome Project (N=45) and the Dunedin Study (N=20) were poor (ICCs=.067-.485). Collectively, these findings demonstrate that common task-fMRI measures are not currently suitable for brain biomarker discovery or individual differences research. We review how this state of affairs came to be and highlight avenues for improving task-fMRI reliability.

## Introduction

Since functional magnetic resonance imaging (fMRI) was introduced in 1992 (Kwong et al., 1992), scientists have had unprecedented ability to non-invasively observe brain activity in behaving humans. In conventional fMRI, regional brain activity is estimated by measuring the blood oxygen level-dependent (BOLD) signal which indexes changes in blood oxygenation associated with neural activity (Logothetis et al., 2001). One of the most common forms of BOLD fMRI is based on tasks during which researchers “map” brain activity associated with specific cognitive functions by contrasting the regional BOLD signal during a control condition with the BOLD signal during a condition of interest. In this way, task-fMRI has given neuroscientists unique insights into the brain basis of human behavior, from basic perception to complex thought, and has given clinicians and mental-health researchers the opportunity to directly measure dysfunction in the organ responsible for disorder.

Originally, task-fMRI was primarily used to understand functions supported by the typical or average human brain by measuring within-subject differences in activation between task and control conditions, and averaging them together across subjects to measure a group effect. To this end, fMRI tasks have been developed and optimized to elicit robust activation in a particular brain region of interest (ROI) or circuit when specific experimental conditions are contrasted. For example, increased amygdala activity is observed when subjects view emotional faces in comparison with geometric shapes and increased ventral striatum activity is observed when subjects win money in comparison to when they lose money (Barch et al., 2013). The robust brain activity elicited using this within-subjects approach led researchers to use the same fMRI tasks to study between-subjects differences. The logic behind this strategy is straightforward: if a brain region activates during a task, then individual differences in the magnitude of that activation may contribute to individual differences in behavior as well as any associated risk for disorder. Thus, if the amygdala is activated when people view threatening stimuli, then differences between people in the degree of amygdala activation should signal differences between them in threat sensitivity and related clinical phenomenon like anxiety and depression (Swartz et al., 2015). In this way, fMRI was transformed from a tool for understanding how the average brain works to a tool for studying how the brains of individuals differ.

The use of task-fMRI to study differences between people heralded the possibility that it could offer a powerful tool for discovering biomarkers for brain disorders (Woo et al., 2017). Broadly, a biomarker is a biological indicator often used for risk stratification, diagnosis, prognosis and evaluation of treatment response. However, to be useful as a biomarker, an indicator must first be reliable. Reliability is the ability of a measure to give consistent results under similar circumstances. It puts a limit on the predictive utility, power, and validity of any measure (see **Box 1** and **Fig. 1**). In this way, reliability is critical for both clinical applications and research practice. Measures with low reliability are unsuitable as biomarkers and cannot predict clinical health outcomes. That is, if a measure is going to be used by clinicians to predict the likelihood that a patient will develop an illness in the future, then the patient cannot score randomly high on the measure at one assessment and low on the measure at the next assessment.

**Fig. 1.**
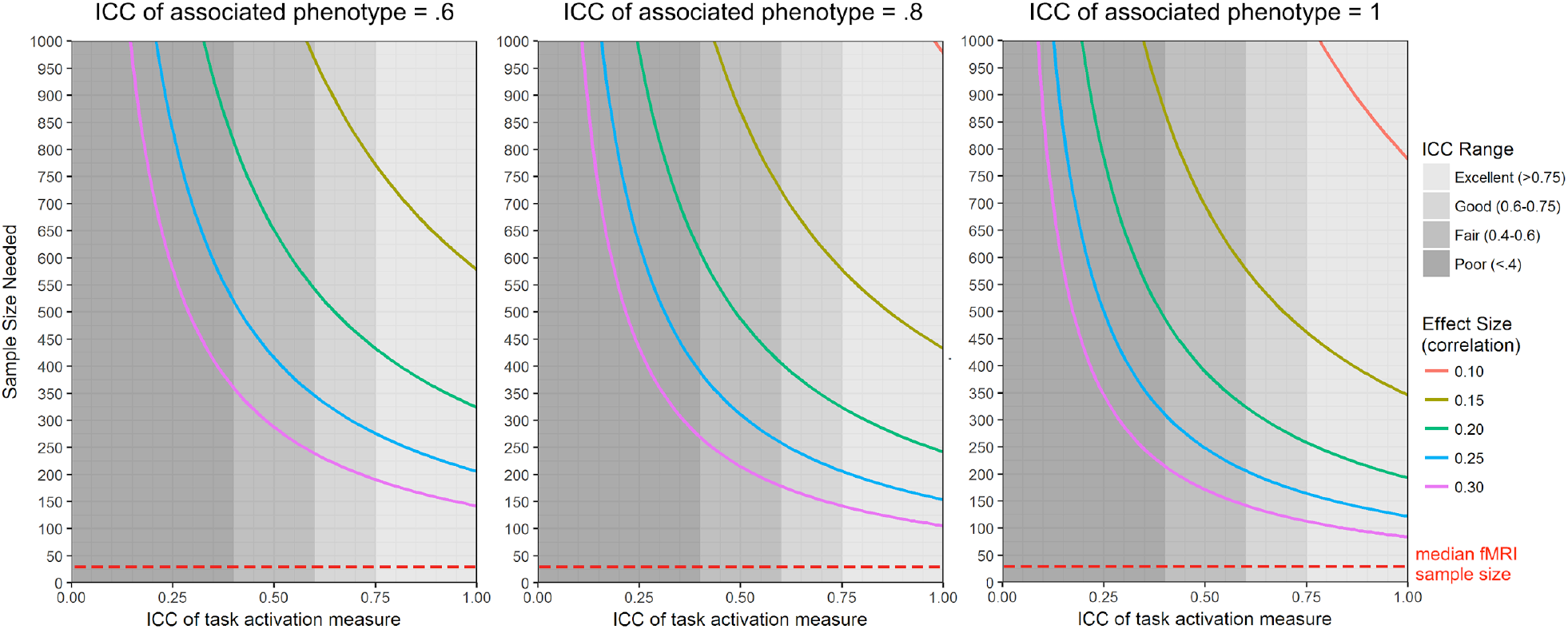
The influence of task-fMRI test-retest reliability on sample size required for 80% power to detect brain-behavior correlations of effect sizes commonly found in psychological research. Power curves are calculated for three levels of reliability of the associated behavioral/clinical phenotype. The figure was generated using the “pwr.r.test” function in R, with the value for “r” specified according to the attenuation formula in Box 1. The figure emphasizes the impact of low reliability at the lower N range because most fMRI studies are relatively small (median N = 28.5 (Poldrack et al., 2017)).

To progress toward a cumulative neuroscience of individual differences with clinical relevance we must establish reliable brain measures. While the reliability of task-fMRI has previously been discussed (Bennett & Miller, 2010; Herting et al., 2018), individual studies provide highly variable estimates, often come from small test-retest samples employing a wide-variety of analytic methods, and sometimes reach-contradictory conclusions about the reliability of the same tasks (Manuck et al., 2007; Nord et al., 2017). This leaves the overall reliability of task-fMRI, as well as the specific reliabilities of many of the most commonly used fMRI tasks, largely unknown. An up-to-date, comprehensive review and meta-analysis of the reliability of task-fMRI and an in-depth examination of the reliability of the most widely used task-fMRI measures is needed. Here, we present evidence from two lines of analysis that point to the poor reliability of commonly used task-fMRI measures. First, we conducted a meta-analysis of the test-retest reliability of regional activation in task-fMRI. Second, in two recently collected datasets, we conducted pre-registered analyses (https://sites.google.com/site/moffittcaspiprojects/home/projectlist/knodt_2019) of the test-retest reliability of brain activation in *a priori* regions of interest across several commonly used fMRI tasks.

## Methods

### Meta-analytic Reliability of Task-fMRI

We performed a systematic review and meta-analysis following PRISMA guidelines (see Supplemental Fig. S1). We searched Google Scholar for peer reviewed articles written in English and published on or before April 1, 2019 that included test-retest reliability estimates of task-fMRI activation. We used the advanced search tool to find articles that include all of the terms “ICC,” “fmri,” and “retest”, and at least one of the terms “ROI,” “ROIs,” “region of interest,” or “regions of interest.” This search yielded 1,170 articles.

#### Study Selection and Data Extraction

One author (MLM) screened all titles and abstracts before the full texts were reviewed (by authors MLE and ARK). We included all original, peer-reviewed empirical articles that reported test-retest reliability estimates for activation during a BOLD fMRI task. All ICCs reported in the main text and supplement were eligible for inclusion. If ICCs were only depicted graphically (e.g. bar graph), we did our best at judging the value from the graph. Voxel-wise ICCs that were only depicted on brain maps were not included. For ICCs calculated based on more than 2 time points, we used the average of the intervals as the value for interval (e.g. the average of the time between time points 1 and 2 and time points 2 and 3 for an ICC based on 3 time points). For articles that reported ICCs from sensitivity analyses in addition to primary analyses on the same data (e.g. using different modeling strategies or excluding certain subjects) we only included ICCs from the primary analysis. We did not include ICCs from combinations of tasks. ICCs were excluded if they were from a longitudinal or intervention study that was designed to assess change, if they did not report ICCs based on measurements from the same MRI scanner and/or task, or if they reported reliability on something other than activation measures across subjects (e.g., spatial extent of activation or multi-voxel patterns of activation within subjects).

Two authors (MLE and ARK) extracted data about sample characteristics (publication year, sample size, healthy versus clinical), study design (test-retest interval, event-related or blocked, task length, and task type), and ICC reporting (i.e., was the ICC thresholded?). For each article, every reported ICC meeting the above study-selection requirements was recorded.

#### Statistical Analyses

For most of the studies included, no standard error or confidence interval for the ICC was reported. Therefore, in order to include as many estimates as possible in the meta-analysis, the standard error of all ICCs was estimated using the Fisher r-to-Z transformation for ICC values (Chen et al., 2018; McGraw & Wong, 1996).

A random-effects multilevel meta-analytic model was fit using tools from the metafor package in R (“Metafor Package R Code for Meta-Analysis Examples,” 2019). In this model, ICCs and standard errors were averaged within each unique sample, task, and test-retest interval (or “substudy”) within each article (or “study”; (Borenstein et al., 2009)). For the results reported in the Main Article, the correlation between ICCs in each substudy was assumed to be 1 so as to ensure that the meta-analytic weight for each substudy was based solely on sample size rather than the number of ICCs reported. However, sensitivity analyses revealed that this decision had very little impact on the overall result (see Supplemental Fig. S2). In the meta-analytic model, substudies were nested within studies to account for the non-independence of ICCs estimated within the same study. Meta-analytic summaries were estimated separately for substudies that reported ICC values that had been thresholded (i.e., when studies calculated multiple ICCs, but only reported values above a minimum threshold) because of the documented spurious inflation of effect sizes that occur when only statistically significant estimates are reported (Kriegeskorte et al., 2009; Poldrack et al., 2017; Vul et al., 2009; Yarkoni, 2009).

To test for effects of moderators, a separate random-effects multilevel model was fit to all 1,146 ICCs (i.e., without averaging within each substudy, since many substudies included ICCs with different values for one or more moderators). The moderators included were task length, task design (block vs event-related), task type (e.g. emotion, executive control, reward, etc), ROI type (e.g. structural or functional), ROI location (cortical vs subcortical), sample type (healthy vs clinical), retest interval, number of citations per year, and whether ICCs were thresholded on significance (see Supplemental Table S1 for descriptive statistics on all moderators tested). All moderators were simultaneously entered into the model as random effects. In the multi-level model, ICCs were nested within substudies, which were in turn nested within studies. This was done to account for the non-independence of ICCs estimated within the same substudy, as well as the non-independence of substudies conducted within the same study.

### Analyses of New Datasets

#### Human Connectome Project (HCP)

This is a publicly available dataset that includes 1,206 participants with extensive structural and functional MRI (Van Essen et al., 2013). In addition, 45 participants completed the entire scan protocol a second time (with a mean interval between scans of approximately 140 days). All participants were free of current psychiatric or neurologic illness and were between 25 and 35 years of age.

The seven tasks employed in the HCP were designed to identify functionally relevant “nodes” in the brain. These tasks included an “n-back” working memory / executive function task (targeting the dorsolateral prefrontal cortex, or dlPFC (Drobyshevsky et al., 2006)), a “gambling” reward / incentive processing task (targeting the ventral striatum (Delgado et al., 2000)), a motor mapping task consisting of foot, hand, and tongue movements (targeting the motor cortex (Drobyshevsky et al., 2006)), an auditory language task (targeting the anterior temporal lobe (Binder et al., 2011)), a social cognition / theory of mind task (targeting the lateral fusiform gyrus, superior temporal sulcus, and other “social-network” regions (Wheatley et al., 2007)), a relational processing / dimensional change detection task (targeting the rostrolateral prefrontal cortex (R. Smith et al., 2007), or rlPFC), and a face-matching emotion processing task (targeting the amygdala (Hariri et al., 2002)).

#### Dunedin Multidisciplinary Health and Development Study

The Dunedin Study is a longitudinal investigation of health and behavior in a complete birth cohort of 1,037 individuals (91% of eligible births; 52% male) born between April 1972 and March 1973 in Dunedin, New Zealand (NZ) and followed to age 45 years (Poulton et al., 2015). Structural and functional neuroimaging data were collected between August 2016 and April 2019, when participants were 45 years old. In addition, 20 Study members completed the entire scan protocol a second time (with a mean interval between scans of 79 days).

Functional MRI was collected during four tasks targeting neural “hubs” in four different domains: a face-matching emotion processing task (targeting the amygdala (Hariri et al., 2002)), a Stroop executive function task (targeting the dlPFC and the dorsal anterior cingulate cortex (Peterson et al., 1999)), a monetary incentive delay reward task (targeting the ventral striatum (Knutson et al., 2000)), and a face-name encoding episodic memory task (targeting the hippocampus (Zeineh et al., 2003)). See Supplemental Methods for additional details, including fMRI pre-processing, for both datasets.

#### ROI Definition

Individual estimates of regional brain activity were extracted according to two commonly used approaches. First, we extracted average values from *a priori* anatomically defined regions. We identified the primary region of interest (ROI) for each task and extracted average BOLD signal change estimates from all voxels within a corresponding bilateral anatomical mask.

Second, we used functionally defined regions based on group-level activation. Here, we generated functional ROIs by drawing 5mm spheres around the group-level peak voxel within the target anatomical ROI for each task (across all subjects and sessions). This is a commonly used strategy for capturing the location of peak activation in each subject despite inter-subject variability in the exact location of the activation. See Supplemental Materials for further details on ROI definition, overlays on the anatomical template (Fig. S3), and peak voxel location (Table S2). We report analyses based on anatomically defined ROIs in the Main Article and report sensitivity analyses using functional ROIs in the Supplement.

#### Reliability Analysis

Subject-level BOLD signal change estimates were extracted for each task, ROI, and scanning session. Reliability was quantified using a 2-way mixed effects intraclass correlation coefficient (ICC), with session modeled as a fixed effect, subject as a random effect, and test-retest interval as an effect of no interest. This mixed effects model is referred to as ICC (3,1) by Shrout and Fleiss (1979), and defined as:

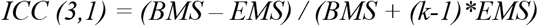

where BMS = between-subjects mean square, EMS = error mean square, and k = number of “raters,” or scanning sessions (in this case 2). We note that ICC (3,1) tracks the consistency of measures between sessions rather than absolute agreement, and is commonly used in studies of task-fMRI test-retest reliability due to the possibility of habituation to the stimuli over time (Plichta et al., 2012).

To test reliability for each task more generally, we calculated ICCs for all target ROIs across all 11 tasks. Since three of the tasks (the emotion, reward, and executive function tasks) were very similar across the HCP and Dunedin Studies and targeted the same region, the same ROI was used for these tasks in both studies, resulting in a total of eight unique target ROIs assessed for reliability. To further visualize global patterns of reliability, we also calculated voxel-wise maps of ICC (3,1) using AFNI’s 3dICC_REML.R function (Chen et al., 2013). Finally, to provide a benchmark for evaluating task-fMRI reliability, we determined the test-retest reliability of three commonly used structural MRI measures: cortical thickness and surface area for each of 360 parcels or ROIs (Glasser et al., 2016) as well as subcortical volume for 17 structures. These analyses were pre-registered (https://sites.google.com/site/moffittcaspiprojects/home/projectlist/knodt_2019). Code and data for this manuscript is available at github.com/HaririLab/Publications/tree/master/ElliottKnodt2020PS_tfMRIReliability

## Results

### Reliability of Individual Differences in Task-fMRI: A Systematic Review and Meta-analysis

We identified 56 articles meeting criteria for inclusion in the meta-analysis, yielding 1,146 ICC estimates derived from 1,088 unique participants across 90 distinct substudies employing 66 different task-fMRI paradigms (**Fig. 2**). These articles were cited a total of 2,686 times, with an average of 48 citations per article and 5.7 citations per article, per year. During the study-selection process, we discovered that some analyses calculated many different ICCs (across multiple ROIs, contrasts, and tasks), but only reported a subset of the estimated ICCs that were either statistically significant or reached a minimum ICC threshold. This practice leads to inflated reliability estimates (Kriegeskorte et al., 2010, 2009; Poldrack et al., 2017). Therefore, we performed separate analyses of data from un-thresholded and thresholded reports.

**Fig. 2.**
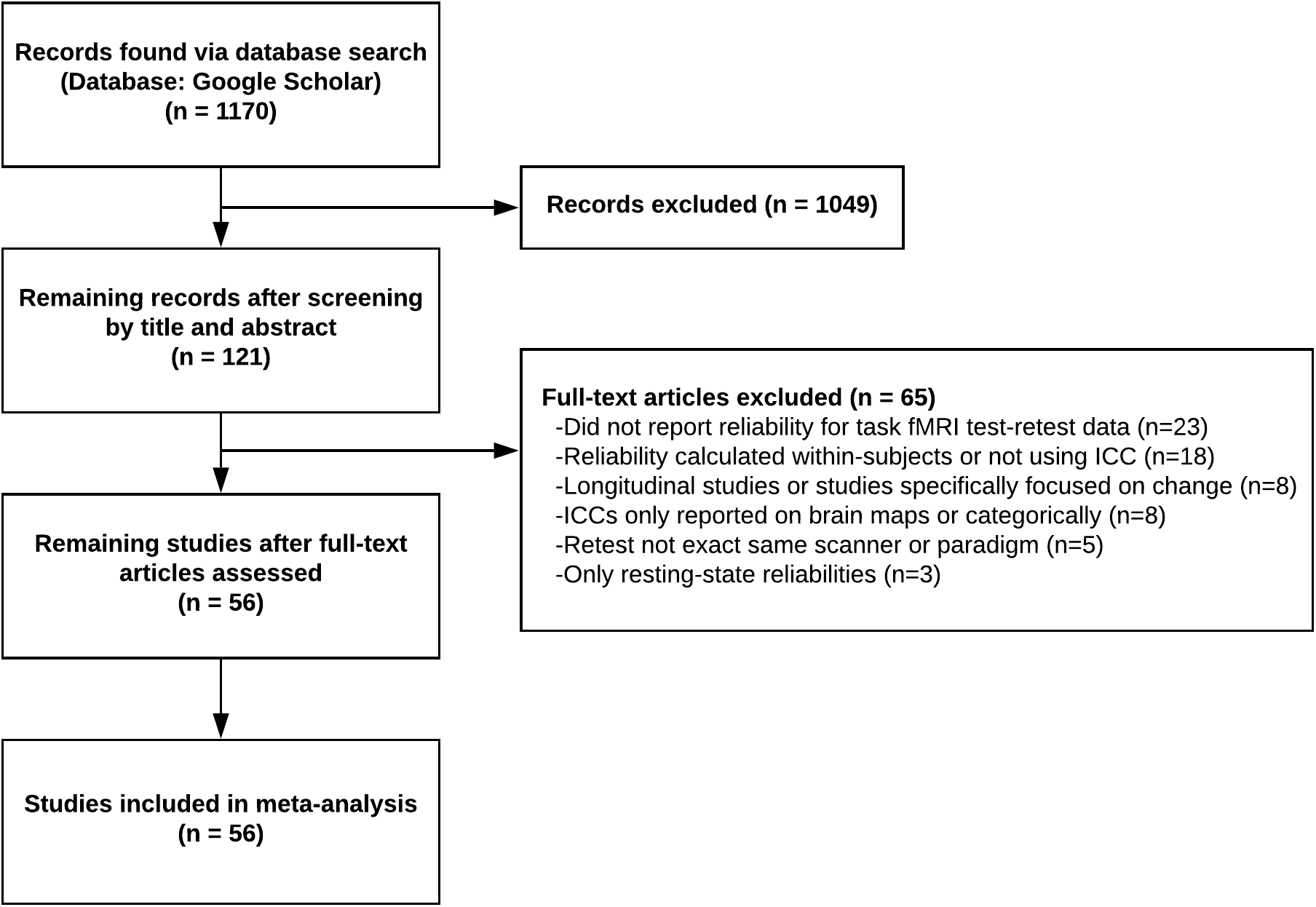
Flow diagram for systematic literature review and meta-analysis.

**Fig. 3** shows the test-retest reliability coefficients (ICCs) from 77 substudies reporting un-thresholded values (average N = 19.6, median N = 17). 56% of the values fell into the range of what is considered “poor” reliability (below .4), an additional 24% of the values fell into the range of what is considered “fair” reliability (.4 - .6), and only 20% fell into the range of what is considered “good” (.6 - .75) or “excellent” (above .75) reliability. A random effects meta-analysis revealed an average ICC of .397 (95% CI, .330 - .460; P < .001), which is in the “poor” range (Cicchetti & Sparrow, 1981). There was evidence of between-study heterogeneity (I^2^ = 31.6; P = 0.04).

**Fig. 3.**
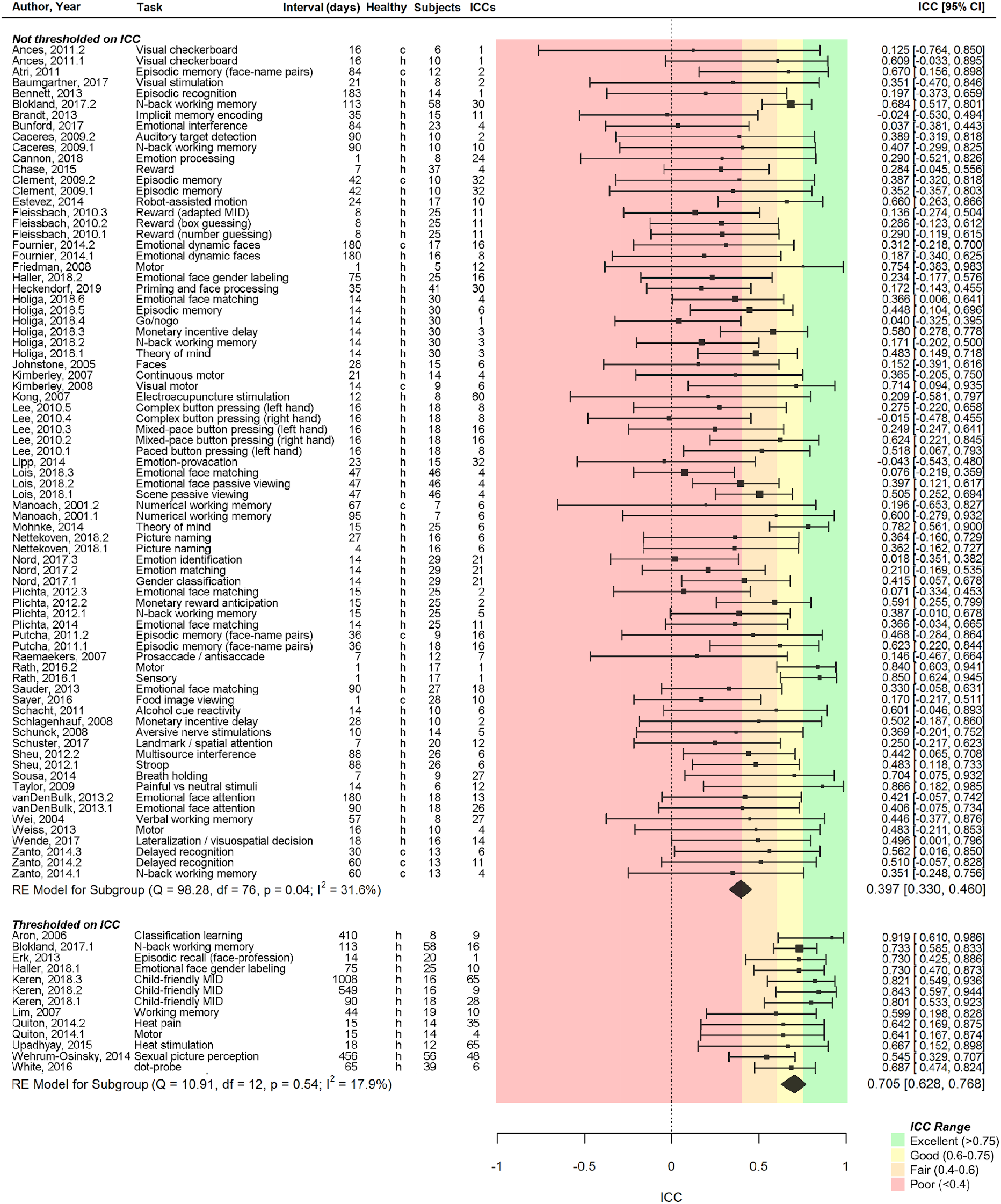
Forest plot for the results of the meta-analysis of task-fMRI test-retest reliability. The forest plot displays the estimate of test-retest reliability of each task-fMRI measure from all ICCs reported in each study. Each substudy is labelled as h if the sample in the study consisted of healthy controls or c if the study consisted of a clinical sample. Studies are split into two sub-groups. The first group of studies reported all ICCs that were calculated, thereby allowing for a relatively unbiased estimate of reliability. The second group of studies selected a subset of calculated ICCs based on the magnitude of the ICC or another non-independent statistic, and then only reported ICCs from that subset. This practice leads to inflated reliability estimates and therefore these studies were meta-analyzed separately to highlight this bias.

As expected, the meta-analysis of 13 substudies that only reported ICCs above a minimum threshold (average N = 24.2, median N = 18) revealed a higher meta-analytic ICC of .705 (95% CI, .628 - .768; P < .001; I^2^ = 17.9). This estimate, which is 1.78 times the size of the estimate from un-thresholded ICCs, is in the good range, suggesting that the practice of thresholding inflates estimates of reliability in task-fMRI. There was no evidence of between-study heterogeneity (I^2^ = 17.9; P = 0.54).

A moderator analysis of all substudies revealed significantly higher reliability for studies that thresholded based on ICC (QM = 6.531, df = 1, P = .010; β = .140). In addition, ROIs located in the cortex had significantly higher ICCs than those located in the subcortex (QM = 114.476, df = 1, P < .001; β= .259). However, we did not find evidence that the meta-analytic estimate was moderated by task type, task design, task length, test-retest interval, ROI type, sample type, or number of citations per year. Finally, we tested for publication bias using the Egger random effects regression test (Egger et al., 1997) and found no evidence for bias (Z = .707, P = .480).

The results of the meta-analysis were illuminating, but not without interpretive difficulty. First, the reliability estimates came from a wide array of tasks and samples, so a single meta-analytical reliability estimate could obscure truly reliable task-fMRI paradigms. Second, the studies used different (and some, now outdated) scanners and different pre-processing and analysis pipelines, leaving open the possibility that reliability has improved with more advanced technology and consistent practices. To address these limitations and possibilities, we conducted pre-registered analyses of two new datasets, using state-of-the-art scanners and practices to assess individual differences in commonly used tasks tapping a variety of cognitive and affective functions.

### Reliability of Individual Differences in Task-fMRI: Pre-registered Analyses in Two New Datasets

We evaluated test-retest reliabilities of activation in *a priori* regions of interest for 11 commonly used fMRI tasks (see **Methods**). In the Human Connectome Project (HCP), 45 participants were scanned twice using a custom 3T Siemens scanner, on average 140 days apart (sd = 67.1 days), using seven tasks targeting emotion, reward, executive function, motor, language, social cognition, and relational processing. This sample size was determined by the publicly available data in the HCP. In the Dunedin Study, 20 participants were scanned twice using a 3T Siemens Skyra, on average 79 days apart (sd = 10.3 days), using four tasks targeting emotion, reward, executive control, and episodic memory. This sample size corresponds to the average sample size used in the meta-analyzed studies. Three of the tasks were similar across the two studies, allowing us to test the replicability of task-fMRI reliabilities. For each of the eight unique tasks across the two studies, we identified the task’s primary target region, resulting in a total of eight *a priori* ROIs (see **Methods**).

#### Group-level activation

To ensure that the 11 tasks were implemented and processed correctly, we calculated the group-level activation in the target ROIs using the primary contrast of interest for each task (see Supplemental Methods for details). These analyses revealed that each task elicited the expected robust activation in the target ROI at the group level (i.e., across all subjects and sessions; see warm-colored maps in **Fig. 4** for the three tasks in common between the two studies and Supplemental Fig. S4 for remaining tasks).

**Fig. 4.**
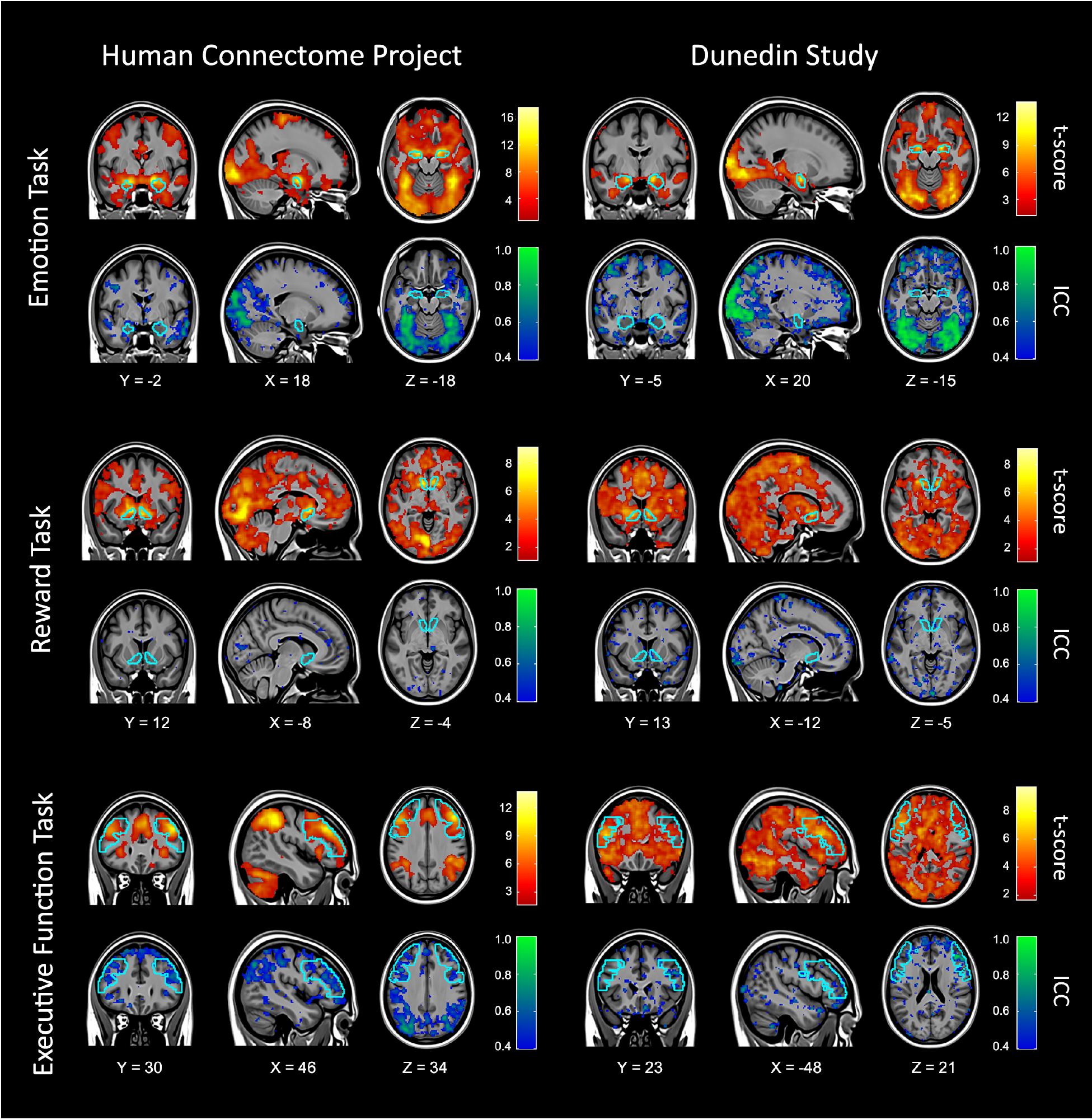
Whole-brain activation and reliability maps for three task-fMRI measures used in both the Human Connectome Project and Dunedin Study. For each task, a whole-brain activation map of the primary within-subject contrast (t-score) is displayed in warm colors (top) and a whole-brain map of the between-subjects reliability (ICC) is shown in cool colors (bottom). For each task, the target ROI is outlined in sky-blue. The activation maps are thresholded at p < .05 whole-brain corrected for multiple comparisons using threshold-free cluster enhancement (Smith & Nichols, 2009). The ICC maps are thresholded so that voxels with ICC < .4 are not colored. These images illustrate that despite robust within-subjects whole-brain activation produced by each task, there is poor between-subjects reliability in this activation, not only in the target ROI but across the whole-brain.

#### Reliability of regional activation

We investigated the reliability of task activation in both datasets using four steps. First, we tested the reliability of activation in the target ROI for each task. Second, for each task we also evaluated the reliability of activation in the other seven *a priori* ROIs. This was done to test if the reliability of target ROIs was higher than the reliability of activation in other (“non-target”) brain regions and to identify any tasks or regions with consistently high reliability. Third, we re-estimated reliability using activation in the left and right hemispheres separately to test if the estimated reliability was harmed by averaging across the hemispheres. Fourth, we tested if the reliability depended on whether ROIs were defined structurally (i.e., using an anatomical atlas) or functionally (i.e., using a set of voxels based on the location of peak activity). See Supplemental Fig. S5 for ICCs of behavior during each fMRI task.

#### Reliability of regional activation in the Human Connectome Project

First, as shown by the estimates circled in black in **Fig. 5**, across the seven fMRI tasks, activation in anatomically defined target ROIs had low reliability (mean ICC = .251; 95% CI, .142 - .360). Only the language processing task had greater than “poor” reliability (ICC = .485). None of the reliabilities entered the “good” range (ICC > .6).

**Fig. 5.**
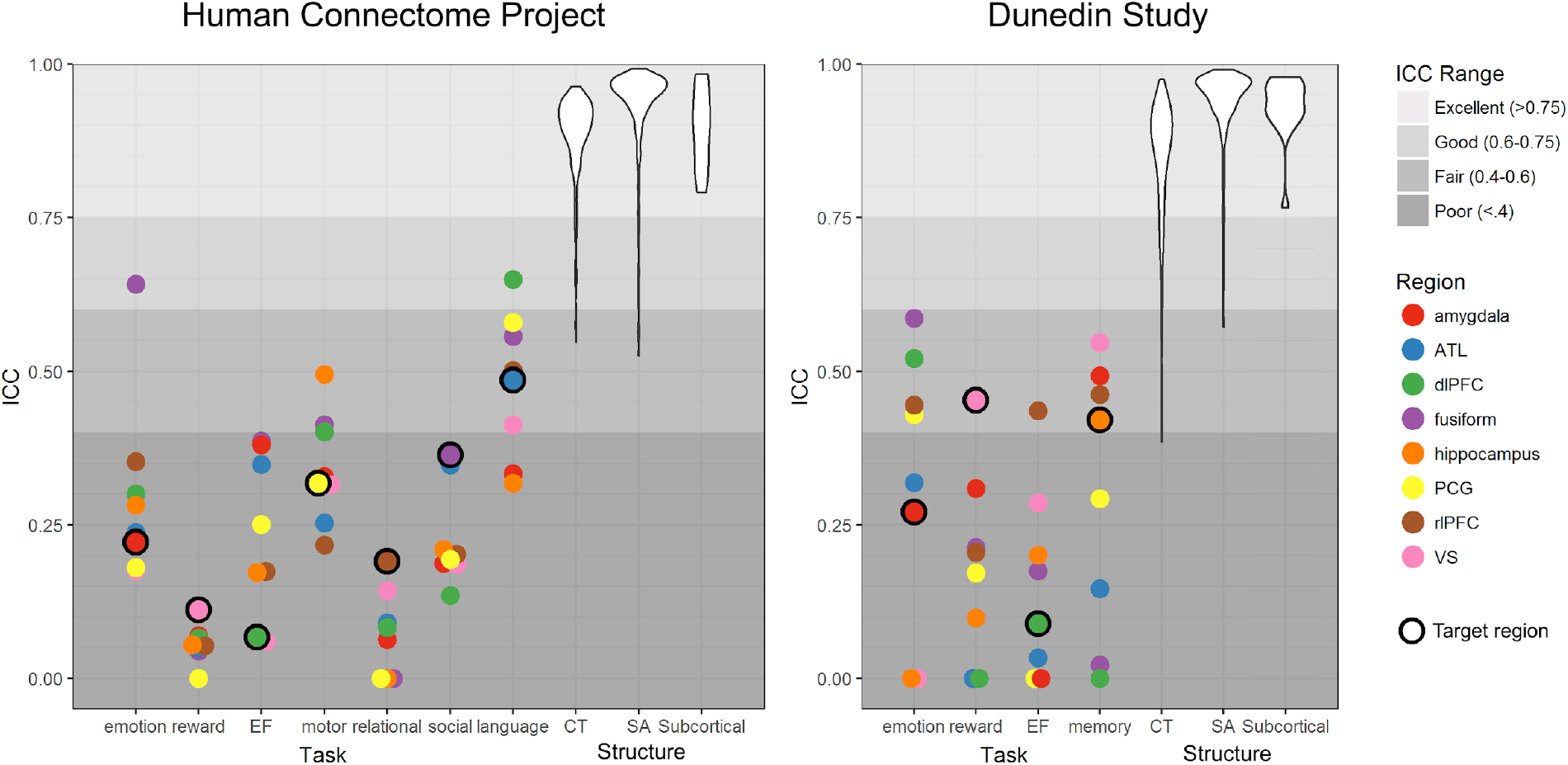
Test-retest reliabilities of region-wise activation measures in 11 commonly used task-fMRI paradigms (EF = executive function). For each task, ICCs were estimated for activation in the *a priori* target ROI (circled in black) and non-target ROIs selected from the other tasks. These plots show that task-fMRI measures of regional activation in both the Human Connectome Project and Dunedin Study are generally unreliable and the ROIs that are “targeted” by the task are rarely more reliable than non-target ROIs (ATL = anterior temporal lobe, dlPFC = dorsolateral prefrontal cortex, PCG = precentral gyrus, rlPFC = rostrolateral prefrontal cortex, VS = ventral striatum). As a benchmark, ICCs of three common structural MRI measures (CT = Cortical Thickness, SA = Surface Area, and Subcortical Volume) are depicted as violin plots representing the distribution of ICCs for each of the 360 parcels for CT and SA, and 17 subcortical structures for grey matter volume. Note that negative ICCs are set to 0 for visualization.

Second, the reliability of task activation in non-target ROIs was also low (**Fig. 5**; mean ICC = .239; 95% CI, .188 - .289), but not significantly lower than the reliability in target ROIs (P = .474).

Third, the reliability of task activation calculated from left and right ROIs separately resembled estimates from averaged ROIs (mean left ICC = .207 in target ROIs and .196 in non-target ROIs, mean right ICC = .259 in target ROIs and .236 in non-target ROIs; Supplemental Fig. S6).

Fourth, the reliability of task activation in functionally defined ROIs was also low (mean ICC = .381; 95% CI, .317 - .446), with only the motor and social tasks exhibiting ICCs greater than .4 (ICCs = .550 and .446 respectively; see Supplemental Fig. S6).

As an additional step, to account for the family structure present in the HCP, we re-estimated reliability after removing one of each sibling/twin pair in the test-retest sample. Reliability in bilateral anatomical ROIs in the subsample of N=26 unrelated individuals yielded reliabilities very similar to the overall sample (mean ICC = .301 in target ROIs and .218 in non-target ROIs; Supplemental Fig. S6).

#### Reliability of regional activation in the Dunedin Study

First, as shown by the estimates circled in black in **Fig. 5**, activation in the anatomically defined target ROI for each of the four tasks had low reliability (mean ICC = .309; 95% CI, .145 - .472), with no ICCs reaching the “good” range (ICC > .6).

Second, the reliability of activation in the non-target ROIs was also low (**Fig. 5**; mean ICC = .193; 95% CI, .100 - .286), but not significantly lower than the reliability in target ROIs (P = .140).

Third, the reliability of task activation calculated for the left and right hemispheres separately was similar to averaged ROIs (mean left ICC = .243 in target ROIs and .202 in non-target ROIs, mean right ICC = .358 in target ROIs and .192 in non-target ROIs; Supplemental Fig. S6).

Fourth, functionally defined ROIs again did not meaningfully improve reliability (mean ICC = .325; 95% CI, .197 - .453; see Supplemental Fig. S6).

#### Reliability of structural measures

To provide a benchmark for evaluating the test-retest reliability of task-fMRI, we investigated the reliability of three commonly used structural MRI measures: cortical thickness, surface area and subcortical grey matter volume. Consistent with prior evidence (Han et al., 2006; Maclaren et al., 2014) that structural MRI phenotypes have excellent reliability (i.e., ICCs > .9), global and regional structural MRI measures in the present samples demonstrated very high test-retest reliabilities (**Fig. 5**). For average cortical thickness, ICCs were .953 and .939 in the HCP and Dunedin Study datasets, respectively. In the HCP, parcel-wise (i.e., regional) cortical thickness reliabilities averaged .886 (range .547 - .964), with 100% crossing the “fair” threshold, 98.6% the “good” threshold, and 94.2% the “excellent” threshold. In the Dunedin Study, parcel-wise cortical thickness reliabilities averaged .846 (range .385 - .975), with 99.7% of ICCs above the “fair” threshold, 96.4% above “good”, and 84.7% above “excellent.” For total surface area, ICCs were .999 and .996 in the HCP and Dunedin Study datasets, respectively. In the HCP, parcel-wise surface area ICCs averaged .937 (range .526 - .992), with 100% crossing the “fair” threshold, 98.9% crossing the “good” threshold, and 96.9% crossing the “excellent” threshold. In the Dunedin Study, surface area ICCs averaged .942 (range .572 - .991), with 100% above the “fair” threshold, 99.7% above “good,” and 98.1% above “excellent.” For subcortical volumes, ICCs in the HCP averaged .903 (range .791 - .984), with all ICCs above the “excellent” threshold. In the Dunedin Study, subcortical volumes averaged .931 (range .767 - .979), with all ICCs above the “excellent” threshold. See Supplemental Table S3 for reliabilities of each subcortical region evaluated.

## Discussion

We found evidence that commonly used task-fMRI measures generally do not have the test-retest reliability necessary for biomarker discovery or brain-behavior mapping. Our meta-analysis of task-fMRI reliability revealed an average test-retest reliability coefficient of .397, which is below the minimum required for good reliability (ICC = .6 (Cicchetti & Sparrow, 1981)) and far below the recommended cutoffs for clinical application (ICC = .8) or individual-level interpretation (ICC = .9) (Guilford, 1946). Of course, not all task-fMRI measures are the same, and it is not possible to assign a single reliability estimate to all individual-difference measures gathered in fMRI research. However, we found little evidence that task type, task length, or test-retest interval had an appreciable impact on the reliability of task-fMRI.

We additionally evaluated the reliability of 11 commonly used task-fMRI measures in the HCP and Dunedin Study. Unlike many of the studies included in our meta-analysis, these two studies were completed recently on modern scanners using cutting-edge acquisition parameters, up-to-date artifact reduction, and state-of-the-art preprocessing pipelines. Regardless, the average test-retest reliability was again poor (ICC = .228). In these analyses, we found no evidence that ROIs “targeted” by the task were more reliable than other, non-target ROIs (mean ICC = .270 for target, .228 for non-target) or that any specific task or target ROI consistently produced measures with high reliability. Of interest, the reliability estimate from these two studies was considerably smaller than the meta-analysis estimate (meta-analytic ICC = .397), possibly due to the phenomenon that pre-registered analyses often yield smaller effect sizes than analyses from publications without pre-registration, which affords increased flexibility in analytic decision-making (Schäfer & Schwarz, 2019).

### The two disciplines of fMRI research

Our results harken back to Lee Cronbach’s classic 1957 article in which he described the “two disciplines of scientific psychology” (Cronbach, 1957). According to Cronbach, the “experimental” discipline strives to uncover universal human traits and abilities through experimental control and group averaging, whereas the “correlational” discipline strives to explain variation between people by measuring how they differ from one another. A fundamental distinction between the two disciplines is how they treat individual differences. For the experimental researcher, variation between people is error that must be minimized to detect the largest experimental effect. For the correlational investigator, variation between people is the primary unit of analysis and must be measured carefully to extract reliable individual differences (Cronbach, 1957; Hedge et al., 2018).

Current task-fMRI paradigms are largely descended from the “experimental” discipline. Task-fMRI paradigms are intentionally designed to reveal how the average human brain responds to provocation, while minimizing between-subject variance. Paradigms that are able to elicit robust targeted brain activity at the group-level are subsequently converted into tools for assessing individual differences. Within-subject robustness is, then, often inappropriately invoked to suggest between-subject reliability, despite the fact that reliable within-subject experimental effects at a group level can arise from unreliable between-subjects measurements (Fröhner et al., 2019).

This reasoning is not unique to task-fMRI research. Behavioral measures that elicit robust within-subject (i.e., group) effects have been shown to have low between-subjects reliability; for example, the mean test-retest reliability of the Stroop Test (ICC = .45; (Hedge et al., 2018)) is strikingly similar to the mean reliability of our task-fMRI meta-analysis (ICC = .397). Nor is it the case that MRI measures, or even the BOLD signal itself, are inherently unreliable. Both structural MRI measures in our analyses (see Fig. 5), as well as measures of intrinsic functional connectivity estimated from long fMRI scans (Elliott et al., 2019; Gratton et al., 2018), demonstrate high test-retest reliability. Thus, it is not the tool that is problematic but rather the strategy of adopting tasks developed for experimental cognitive neuroscience that appear to be poorly suited for reliably measuring differences in brain activation between people.

### Recommendations and Future Directions

We next consider several avenues for maximizing the value of existing datasets as well as improving the reliability of task-fMRI moving forward. We begin with recommendations that can be implemented immediately (1, 2), before moving on to recommendations that will require additional data collection and innovation (3, 4).

#### 1) Immediate opportunities for task-fMRI: from brain hotspots to whole-brain signatures

Currently, the majority of task-fMRI measures are based on contrasts between conditions (i.e., change scores), extracted from ROIs. However, change scores will always have lower reliability than their constituent measures (Hedge et al., 2018), and have been shown to undermine the reliability of task-fMRI (Infantolino et al., 2018). However, contrast-based activation values extracted from ROIs represent only one possible measure of individual differences that can be derived from task-fMRI data. For example, several multivariate methods have been proposed to increase the reliability and predictive utility of task-fMRI measures by exploiting the high dimensionality inherent in fMRI data (Dubois & Adolphs, 2016; Yarkoni & Westfall, 2017). To name a few, the reliability of task-fMRI may be improved by developing measures with latent variable models (Cooper et al., 2019), measuring individual differences in representational spaces with multi-voxel pattern analysis (Norman et al., 2006), and training cross-validated machine learning models that establish reliability through prediction of individual differences in independent samples (Yarkoni & Westfall, 2017). In addition, in many already-collected datasets, task-fMRI can be combined with resting-state fMRI data to produce reliable measures of intrinsic functional connectivity (Elliott et al., 2019; Greene et al., 2018). Thus, there are multiple available approaches to maximizing the value of existing task-fMRI datasets in the context of biomarker discovery and individual differences research.

#### 2) Create a norm of reporting the reliability of task-fMRI measures

The “replicability revolution” in psychological science (Nosek et al., 2015) provides a timely example of how rapidly changing norms can shape research practices and standards. In just a few years, practices to enhance replicability, like pre-registration of hypotheses and analytic strategies, have risen in popularity (Nosek et al., 2018). We believe similar norms would be beneficial for task-fMRI in the context of biomarker discovery and brain-behavior mapping. In particular, researchers should report the reliabilities for all task-fMRI measures whenever they are used to study individual differences (Parsons et al., 2019). In doing so, however, researchers need to ensure adequate power to evaluate test-retest reliability with confidence. Given that correlations begin to stabilize with around 150 observations (Schönbrodt & Perugini, 2013), our confidence in knowing “the” reliability of any specific task will depend on collecting larger test-retest datasets. We provide evidence that the task-fMRI literature generally has low reliability; however, due to the relatively small size of each test-retest sample reported here, we urge readers to avoid making strong conclusions about the reliability of specific fMRI tasks. In the pursuit of precise reliability estimates, it will be important for researchers to collect larger test-retest samples, explore test-retest moderators (e.g. test-retest interval) and avoid reporting inflated reliabilities that can arise from circular statistical analyses (for detailed recommendations see (Kriegeskorte et al., 2010, 2009; Vul et al., 2009)).

Researchers can also provide evidence of between-subjects reliability in the form of internal consistency. While test-retest reliability provides an estimate of stability over time that is suited for trait and biomarker research, it is a conservative estimate that requires extra data collection and can be undermined by habituation effects and rapid fluctuations (Hajcak et al., 2017). In some cases, internal consistency will be more practical because it is cheaper, as it does not require additional data collection and can be used in any situation where the task-fMRI measure of interest is comprised of multiple trials (Streiner, 2003). Internal consistency is particularly well-suited for measures that are expected to change rapidly and index transient psychological states (e.g., current emotions or thoughts). However, internal consistency alone is not adequate for prognostic biomarkers. Establishing a norm of explicitly reporting measurement reliability would increase the replicability of task-fMRI findings and accelerate biomarker discovery.

#### 3) More data from more subjects

Our ability to detect reliable individual differences using task-fMRI will depend, in part, on the field embracing two complementary improvements to the status quo: 1) more subjects per study and 2) more data per subject. It has been suggested that neuroscience is generally an underpowered enterprise, and that small sample sizes undermine fMRI research in particular (Button et al., 2013; Szucs & Ioannidis, 2017). The results presented here suggest that this “power failure” may be further compounded by low reliability in task-fMRI. The median sample size in fMRI research is 28.5 (Poldrack et al., 2017). However, as shown in Fig. 1, task-fMRI measures with ICCs of .397 (the meta-analytic mean reliability) would require N > 214 to achieve 80% power to detect brain-behavior correlations of .3, a moderate effect size equal to the size of the largest replicated brain-behavior associations (Elliott et al., 2018; Nave et al., 2019). For r = .1 (a small effect size common in psychological research (Funder & Ozer, 2019)), adequately powered studies require N > 2,000. And, these calculations are actually best-case scenarios given that they assume perfect reliability of the second “behavioral” variable (see Figure 1). Increasing the sample size of task-fMRI studies and requiring power analyses that take into account unreliability represent a meaningful way forward for boosting the replicability of individual differences research with task-fMRI.

Without substantially higher reliability, task-fMRI measures will fail to provide biomarkers that are meaningful on an individual level. One promising method to improve the reliability of fMRI is to collect more data per subject. Increasing the amount of data collected per subject has been shown to improve the reliability of functional connectivity (Elliott et al., 2019; Gratton et al., 2018) and preliminary efforts suggest this may be true for task-fMRI as well (Gordon et al., 2017). Pragmatically, collecting additional fMRI data will be burdensome for participants, especially in children and clinical populations, where longer scan times often result in greater data artifacts particularly from increased motion. Naturalistic fMRI represents one potential solution to this challenge. In naturalistic fMRI, participants watch stimulus-rich movies during scanning instead of completing traditional cognitive neuroscience tasks. Initial efforts suggest that movie watching is highly engaging for subjects, allows more data collection with less motion and may even better elicit individual differences in brain activity by emphasizing ecological validity over experimental control (Vanderwal et al., 2018). As the field launches large-scale neuroimaging studies (e.g. HCP, UK Biobank, ABCD) in the pursuit of brain biomarkers of disease risk, it is critical that we are confident in the psychometric properties of task-fMRI measurements. This will require funders to advocate and support the collection of more data from more subjects.

#### 4) Develop tasks from the ground up to optimize reliable and valid measurement

Instead of continuing to adopt fMRI tasks from experimental studies emphasizing within-subjects effects, we need to develop new tasks (and naturalistic stimuli) from the ground up with the goal of optimizing their utility in individual differences research (i.e., between-subjects effects). Psychometrics provides many tools and methods for developing reliable individual differences measures that have been underutilized in task-fMRI development. For example, stimuli in task-fMRI could be selected based on their ability to maximally distinguish groups of subjects or to elicit reliable between subject variance. As noted in recommendation 1, psychometric tools for test construction could be adopted to optimize reliable task-fMRI measures including item analysis, latent variable modelling, and internal-consistency measures (Crocker & Algina, 2006).

## Conclusion

A prominent goal of task-fMRI research has been to identify abnormal brain activity that could aid in the diagnosis, prognosis, and treatment of brain disorders. We find that commonly used task-fMRI measures lack minimal reliability standards necessary for accomplishing this goal. Intentional design and optimization of task-fMRI paradigms are needed to measure reliable variation between individuals. As task-fMRI research faces the challenges of reproducibility and replicability, we draw attention to the importance of reliability as well. In the age of individualized medicine and precision neuroscience, funding is needed for novel task-fMRI research that embraces the psychometric rigor necessary to generate clinically actionable knowledge.

## Box 1: Why is reliability critical for task-fMRI research?

Test-retest reliability is widely quantified using the intraclass correlation coefficient (ICC (Shrout & Fleiss, 1979)). ICC can be thought of as the proportion of a measure’s total variance that is accounted for by variation between individuals. An ICC can take on values between −1 and 1, with values approaching 1 indicating nearly perfect stability of individual differences across test-retest measurements, and values at or below 0 indicating no stability. Classical test theory states that all measures are made up of a true score plus measurement error (Novick, 1965). The ICC is used to estimate the amount of reliable, true-score variance present in an individual differences measure. When a measure is taken at two timepoints, the variance in scores that is due to measurement error will consist of random noise and will fail to correlate with itself across test-retest measurements. However, the variance in a score that is due to true score will be stable and correlate with itself across timepoints (Crocker & Algina, 2006). Measures with ICC < .40 are thought to have “poor” reliability, those with ICCs between .40 - .60 “fair” reliability, .60 - .75 “good” reliability, and > .75 “excellent” reliability. An ICC > .80 is considered a clinically required standard for reliability in psychology (Cicchetti & Sparrow, 1981).

Reliability is critical for research because the correlation observed between two measures, A and B, is constrained by the square root of the product of each measure’s reliability (Nunnally, 1959):

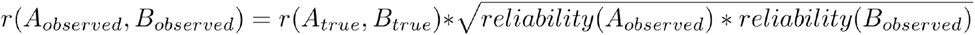

Low reliability of a measure reduces statistical power and increases the sample size required to detect a correlation with another measure. **Fig. 1** shows sample sizes required for 80% power to detect correlations between aa task-fMRI measure of individual differences in brain activation and a behavioral/clinical phenotype, across a range of reliabilities of the task-fMRI measure and expected effect sizes. Power curves are given for three levels of reliability of the hypothetical behavioral/clinical phenotype, where the first two panels (behavioral ICC = .6 and .8) represent most typical scenarios.

## Author Contributions

A.C., A.R.H., T.E.M, M.L.E., and A.R.K. conceived the study and data analysis plan. M.L.E., A.R.K., and M.L.S. prepared MRI data for analysis. M.L.M prepared data for meta-analysis. A.R.K., M.L.E., and M.L.S. conducted the analyses. M.L.E., A.R.K., A.C., A.R.H., and T.E.M. wrote the manuscript. A.C., A.R.H., T.E.M., and R.P. designed, implemented, and/or oversaw the collection and generation of the research protocol. S.R., D.I., and A.R.K. oversaw data collection. All authors discussed the results and contributed to the revision of the manuscript.

## Acknowledgments

Data were provided [in part] by the Human Connectome Project, WU-Minn Consortium (Principal Investigators: David Van Essen and Kamil Ugurbil; 1U54MH091657) funded by the 16 NIH Institutes and Centers that support the NIH Blueprint for Neuroscience Research; and by the McDonnell Center for Systems Neuroscience at Washington University.

The Dunedin Study was approved by the NZ-HDEC (Health and Disability Ethics Committee). The Dunedin Study is supported by NIA grants R01AG049789 and R01AG032282 and U.K. Medical Research Council grant P005918. The Dunedin Multidisciplinary Health and Development Research Unit is supported by the New Zealand Health Research Council and the New Zealand Ministry of Business, Innovation and Employment (MBIE). MLE is supported by the National Science Foundation Graduate Research Fellowship under Grant No. NSF DGE-1644868. Thanks to the members of the Advisory Board for the Dunedin Neuroimaging Study. The authors would also like to thank Tim Strauman and Ryan Bogdan for their feedback on an initial draft of this manuscript, as well as extensive feedback from peer reviewers.

