## supplemental for "What is the test-retest reliability of common task-fMRI measures? New empirical evidence and a meta-analysis"

**This file includes:**

Supplementary Methods
Figs. S1 to S7
Tables S1 to S4
References for SI reference citations

**Supplementary Methods**

**Meta-analysis**

We identified 56 articles meeting the inclusion criteria that were all included in the final meta-analysis (Ances, Vaida, Ellis, & Buxton, 2011; Aron, Gluck, & Poldrack, 2006; Atri et al., 2011; Baumgartner et al., 2017; Bennett & Miller, 2013; Blokland et al., 2017; Brandt et al., 2013; Bunford, Kinney, Michael, & Klumpp, 2017; Caceres, Hall, Zelaya, Williams, & Mehta, 2009; Cannon, Cao, Mathalon, Gee, & on behalf of the NAPLS consortium, 2018; Chase et al., 2015; Clément & Belleville, 2009; Drew Sayer et al., 2016; Erk et al., 2013; Estévez et al., 2014; Fliessbach et al., 2010; Fournier, Chase, Almeida, & Phillips, 2014; Friedman et al., 2008; Haller et al., 2018; Heckendorf, Bakermans-Kranenburg, van Ijzendoorn, & Huffmeijer, 2019; Holiga et al., 2018; Johnstone et al., 2005; Keren et al., 2018; Kimberley, Birkholz, Hancock, VonBank, & Werth, 2008; Kimberley, Khandekar, & Borich, 2008; Kong et al., 2007; Lee et al., 2010; Lim, Choo, & Chee, 2007; Lipp, Murphy, Wise, & Caseras, 2014; Lois, Kirsch, Sandner, Plichta, & Wessa, 2018; Manoach et al., 2001; Mohnke et al., 2014; Nettekoven, Reck, Goldbrunner, Grefkes, & Lucas, 2018; Nord, Gray, Charpentier, Robinson, & Roiser, 2017; Plichta et al., 2014, 2012; Putcha et al., 2011; Quiton, Keaser, Zhuo, Gullapalli, & Greenspan, 2014; Raemaekers et al., 2007; Rath et al., 2016; Sauder, Hajcak, Angstadt, & Phan, 2013; Schacht et al., 2011; Schlagenhauf et al., 2008; Schunck et al., 2008; Schuster et al., 2017; Sheu, Jennings, & Gianaros, 2012; Sousa, Vilela, & Figueiredo, 2014; Taylor & Davis, 2009; Upadhyay et al., 2015; van den Bulk et al., 2013; Wehrum-Osinsky et al., 2014; Weiss et al., 2013; Wei et al., 2004; Wende et al., 2017; White et al., 2016; Zanto, Pa, & Gazzaley, 2014).

### Datasets

***Human Connectome Project (HCP).*** The MRI acquisition parameters and minimal preprocessing of these data have been described extensively elsewhere (Glasser et al., 2013). Briefly, participants underwent extensive MRI measurement that included T1-weighted structural imaging and nearly two hours of fMRI scanning during resting-state and seven tasks. Task-fMRI is described extensively elsewhere (Barch et al., 2013). For our analyses we used the minimally preprocessed data in volumetric Montreal Neurological Institute (MNI) space ("fMRIVolume" pipeline).

Subjects who did not complete a task at one or both time points were removed from analysis for that task. Thus, the final test-retest dataset used in our reliability analyses included 45 subjects for the reward, motor, and executive function tasks; 44 subjects for the social task; 43 subjects for the language task, and 42 subjects for the emotion and relational tasks.

***Dunedin Multidisciplinary Health and Development Study***. The Dunedin Study is a longitudinal investigation of health and behavior in a complete birth cohort of 1,037 individuals (91% of eligible births; 52% male) born between April 1972 and March 1973 in Dunedin, New Zealand (NZ), and eligible based on residence in the province and participation in the first assessment at age 3. The cohort represents the full range of socioeconomic status on NZ's South Island and matches the NZ National Health and Nutrition Survey on key health indicators (e.g., BMI, smoking, GP visits) (Poulton, Moffitt, & Silva, 2015). The cohort is primarily white (93%). Assessments were carried out at birth and ages 3, 5, 7, 9, 11, 13, 15, 18, 21, 26, 32, 38, and 45 years, when 94% of the 997 study members still alive took part, of whom 93% participated in MRI scanning.

The acquisition parameters and task-fMRI measures for the Dunedin Study have been described in detail elsewhere (Elliott et al., 2019). 20 study members completed the entire scan protocol a second time (mean days between scans = 79). These 20 study members made up the Dunedin test-retest sample used in this manuscript. All 20 study members completed all 4 fMRI tasks at both timepoints.

**Image Pre-processing**

Minimal preprocessing was first applied to all data. For the HCP dataset this was done with the HCP minimal preprocessing pipeline (Glasser et al., 2013). This includes correction for B0 distortion, realignment to the single-band reference image to correct for motion, registration to the subject's structural scan, normalization to the 4D mean, brain masking, and non-linear warping to MNI space. All single-band reference images were visually inspected to ensure proper alignment to the anatomical image.

Similar "minimal preprocessing" steps were applied to the Dunedin Study dataset using custom processing scripts. Anatomical images for each subject were skull-stripped, intensity-normalized, and nonlinearly warped to a study-specific average template in the standard stereotactic space of the Montreal Neurological Institute template using the ANTs SyN registration algorithm (Avants, Epstein, Grossman, & Gee, 2008; Klein et al., 2009). Time series images for each subject were despiked, slice-time-corrected, realigned to the first volume in the time series to correct for head motion using AFNI tools (Cox, 1996), corrected for B0 distortions using SPM's fieldmap toolbox (Jezzard & Balaban, 1995), coregistered to the anatomical image using FSL's Boundary Based Registration (Greve & Fischl, 2009), and spatially normalized into MNI space using the non-linear ANTs SyN warp from the anatomical image. All transformations were concatenated so that a single interpolation was performed. The 4D means from each normalized time series were visually inspected to ensure proper alignment to the anatomical template. Following "minimal preprocessing," normalized time series from both datasets were smoothed to minimize noise and residual difference in gyral anatomy with a Gaussian filter, set at 6-mm full-width at half-maximum. Voxel-wise signal intensities were scaled to yield a time series mean of 100 for each voxel, allowing parameter estimates from the general linear model that was subsequently applied to be interpreted as percent signal change (PSC). The AFNI program 3dREMLfit (Cox, 1996) was used to fit a general linear model for first-level task-fMRI data analyses. Linear contrasts employing canonical hemodynamic response functions were used to estimate condition of interest effects for each task, while controlling for low frequency noise using a number of polynomial regressors appropriate for the length of each task. Contrasts of interest for each task were as follows in the HCP: "Faces > Shapes" for the emotion task, "Win > Loss" for the reward task, "2-back > 0-back" for the executive function N-back task, "Motor > Fixation" for the motor task, "Relation > Match" for the relational task, "Mental > Random" for the social task, and "Story > Math" for the language task; and in the Dunedin Study: "Faces > Shapes" for the emotion task, "Gain Anticipation > Neutral Anticipation" for the reward task, "Incongruent > Congruent" for the executive function Stroop task, and "Encoding > Distractor" for the episodic memory task. Volumes exhibiting excessive motion were identified with framewise displacement and standardized DVARS (Nichols, 2017; Power et al., 2014) thresholds specific to each dataset (0.39 mm and 4.9 for the HCP (Burgess et al., 2016) and 0.5 mm and 2.5 for the Dunedin Study, respectively) and subsequently censored from the GLM. To screen for signal dropout, for each subject we calculated the mean temporal SNR for each task within the target region and tested whether this value was an outlier (defined as < 3 standard deviations below the mean across subjects for the task). One subject in the Dunedin Study was identified as an outlier for tSNR in the reward task (ventral striatum); sensitivity analyses conducted without this subject resulted in a lower ICC for the task, leaving the conclusions unchanged.

Additionally, parcel-wise measures of cortical thickness and surface area in both datasets were extracted using the HCP's "PostFreeSurfer" pipeline applied to the T1- and T2-weighted structural scans. Grey matter volumes for 17 regions of interest were extracted separately from the automatic segmentation ("aseg") step of FreeSurfer version 6.0. FreeSurfer version 6.0 was used because the HCP FreeSurfer pipeline was optimized for the cortical surface, resulting in lower-quality segmentation of subcortical volumes in our dataset.

**Anatomical ROI Definition**

We identified the primary target region of interest (ROI) for each task for use in all analyses (see references for each task in the Methods section of the main text). For simplicity, we selected a single target region for the two tasks that are often used to target more than one region: 1) for the HCP social cognition task, we selected the fusiform gyrus over the other regions described in (Wheatley, Milleville, & Martin, 2007) due to the presence of faces in other tasks, and 2) for the Dunedin executive function Stroop task, we selected the dorsolateral prefrontal cortex (dlPFC) over the dorsal anterior cingulate cortex to parallel the HCP executive function task.

Anatomical ROI masks for each ROI were selected based on alignment to the template as well as prior work for the respective task where applicable. We used anatomical definitions from the Automated Anatomical Labeling (AAL) atlas for the hippocampus and the fusiform gyrus (episodic memory and social cognition tasks), the Harvard Oxford Atlas (HOA) for the precentral gyrus and anterior temporal lobe (motor and language tasks; the latter consisted of the combined temporal pole, anterior superior temporal gyrus, anterior middle temporal gyrus, and anterior inferior temporal gyrus regions), the high-resolution amygdala template generated from 168 Human Connectome Project datasets (emotion tasks; Tyszka et al. 2016), and the Oxford-GSK-Imanova structural and connectivity striatal atlases distributed with FSL for the ventral striatum (reward tasks). The dlPFC (executive function task) was defined using the Brodmann Area Atlas provided by Wake Forest University PickAtlas (WFU PickAtlas; www.fmri.wfubmc.edu/downloads), with Brodmann Areas 9 and 46 combined, dilated by 2 voxels, and the medial aspect removed (Tong et al., 2016). The rostrolateral PFC (relational task) was defined as Brodmann Area 10 using the Brodmann Area Atlas provided with MRIcron (https://www.nitrc.org/projects/mricron). For our primary analyses, we combined the anatomical ROIs from the left and right hemisphere into a single bilateral mask. Further, all ROIs were masked to exclude non-grey matter voxels, using the grey matter probability map from the study-specific average template thresholded at 0.25.

**Fig. S1.** PRISMA Checklist for systematic reviews and meta-analyses.

| Section / topic | # | Checklist item | Reported on page # |
|---|---|---|---|
| **TITLE** | | | |
| Title | 1 | Identify the report as a systematic review, meta-analysis, or both. | 1 |
| **ABSTRACT** | | | |
| Structured summary | 2 | Provide a structured summary including, as applicable: background; objectives; data sources; study eligibility criteria, participants, and interventions; study appraisal and synthesis methods; results; limitations; conclusions and implications of key findings; systematic review registration number. | 2 |
| **INTRODUCTION** | | | |
| Rationale | 3 | Describe the rationale for the review in the context of what is already known. | 3-5 |
| Objectives | 4 | Provide an explicit statement of questions being addressed with reference to participants, interventions, comparisons, outcomes, and study design (PICOS). | 4-5 |
| **METHODS** | | | |
| Protocol and registration | 5 | Indicate if a review protocol exists, if and where it can be accessed (e.g., Web address), and, if available, provide registration information including registration number. | 5-6 |
| Eligibility criteria | 6 | Specify study characteristics (e.g., PICOS, length of follow-up) and report characteristics (e.g., years considered, language, publication status) used as criteria for eligibility, giving rationale. | 5-6 |
| Information sources | 7 | Describe all information sources (e.g., databases with dates of coverage, contact with study authors to identify additional studies) in the search and date last searched. | 5-6 |
| Search | 8 | Present full electronic search strategy for at least one database, including any limits used, such that it could be repeated. | 5 |
| Study selection | 9 | State the process for selecting studies (i.e., screening, eligibility, included in systematic review, and, if applicable, included in the meta-analysis). | 5-6 |
| Data collection process | 10 | Describe method of data extraction from reports (e.g., piloted forms, independently, in duplicate) and any processes for obtaining and confirming data from investigators. | 5-6 |
| Data items | 11 | List and define all variables for which data were sought (e.g., PICOS, funding sources) and any assumptions and simplifications made. | 6 |
| Risk of bias in individual studies | 12 | Describe the methods used for assessing risk of bias of individual studies (including specification of whether this was done at the study or outcome level), and how this information is to be used in any data synthesis. | 6-7 |

| | | | |
|---|---|---|---|
| Summary measures | 13 | State the principal summary measures (e.g., risk ratio, difference in means). | 5-6 |
| Synthesis of results | 14 | Describe the methods of handling data and combining results of studies, if done, including measures of consistency (e.g., $I^2$) for each meta-analysis. | 5-7 |
| Risk of bias across studies | 15 | Specify any assessment of risk of bias that may affect the cumulative evidence (e.g., publication bias, selective reporting within studies). | 5-7 |
| Additional analyses | 16 | Describe methods of additional analyses (e.g., sensitivity or subgroup analyses, meta-regression), if done, indicating which were pre-specified. | 6-7, S10 |
| **RESULTS** | | | |
| Study selection | 17 | Give numbers of studies screened, assessed for eligibility, and included in the review, with reasons for exclusions at each stage, ideally with a flow diagram. | Fig. 2 |
| Study characteristics | 18 | For each study, present characteristics for which data were extracted (e.g., study size, PICOS, follow-up period) and provide the citations. | Fig. 3, Table S1 |
| Risk of bias within studies | 19 | Present data on risk of bias of each study and, if available, any outcome level assessment (see item 12). | 13, Fig S2 |
| Results of individual studies | 20 | For all outcomes considered (benefits or harms), present, for each study: (a) simple summary data for each intervention group (b) effect estimates and confidence intervals, ideally with a forest plot. | Fig. 3 |
| Synthesis of results | 21 | Present results of each meta-analysis done, including confidence intervals and measures of consistency. | 13, Fig. 3 |
| Risk of bias across studies | 22 | Present results of any assessment of risk of bias across studies (see Item 15). | Fig. S2 |
| Additional analysis | 23 | Give results of additional analyses, if done (e.g., sensitivity or subgroup analyses, meta-regression [see Item 16]). | 13, Fig. S2 |
| **DISCUSSION** | | | |
| Summary of evidence | 24 | Summarize the main findings including the strength of evidence for each main outcome; consider their relevance to key groups (e.g., healthcare providers, users, and policy makers). | 18-19 |
| Limitations | 25 | Discuss limitations at study and outcome level (e.g., risk of bias), and at review-level (e.g., incomplete retrieval of identified research, reporting bias). | 18-19 |
| Conclusions | 26 | Provide a general interpretation of the results in the context of other evidence, and implications for future research. | 24 |
| **FUNDING** | | | |
| Funding | 27 | Describe sources of funding for the systematic review and other support (e.g., supply of data); role of funders for the systematic review. | 26 |

**Fig. S2.** Sensitivity analyses for the meta-analysis. Here the correlations between ICCs in each substudy were assumed to be 0, representing the extreme case where the ICCs within each substudy were completely independent.

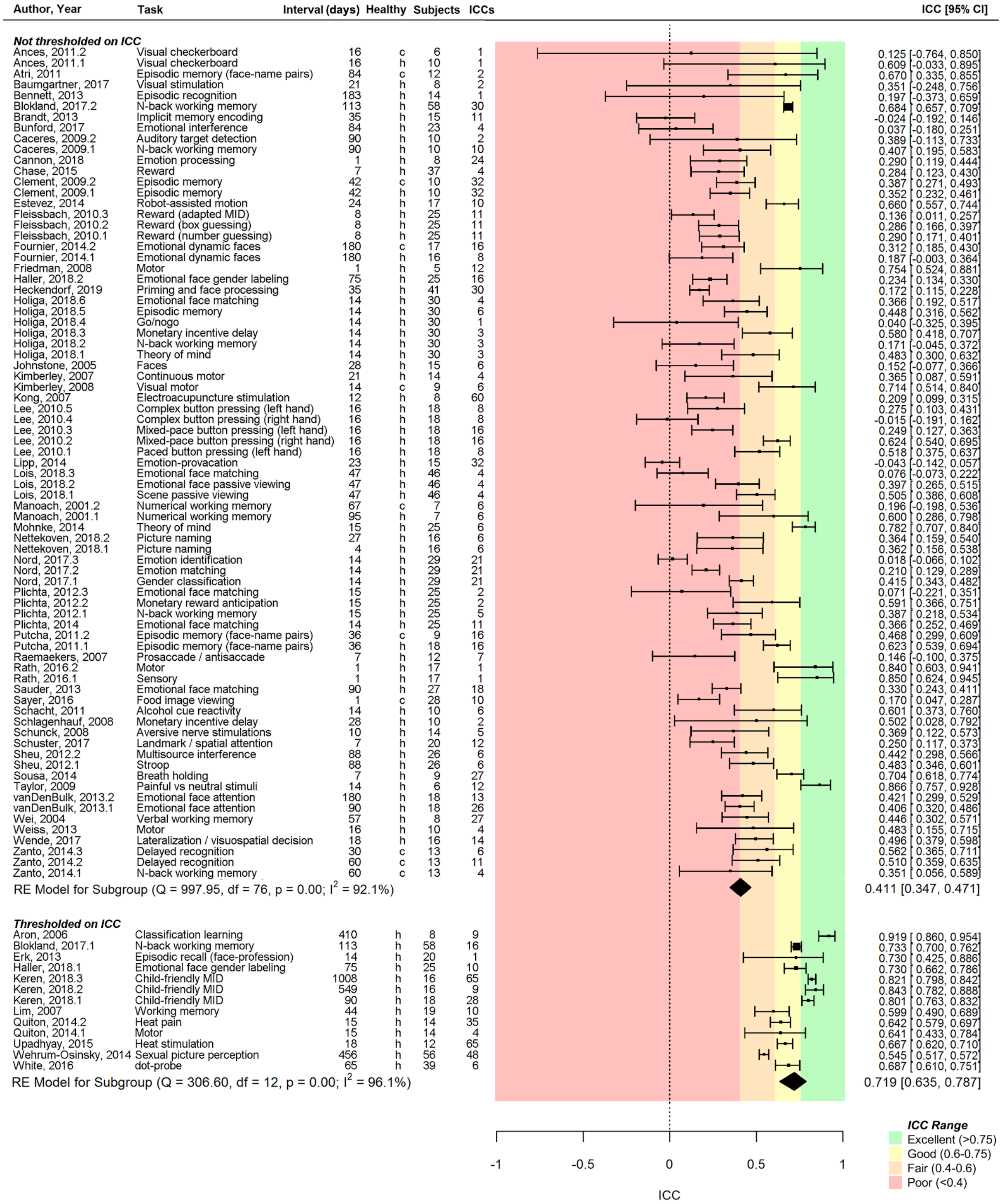

**Fig S3.** ROI masks (red) overlaid on the anatomical template.

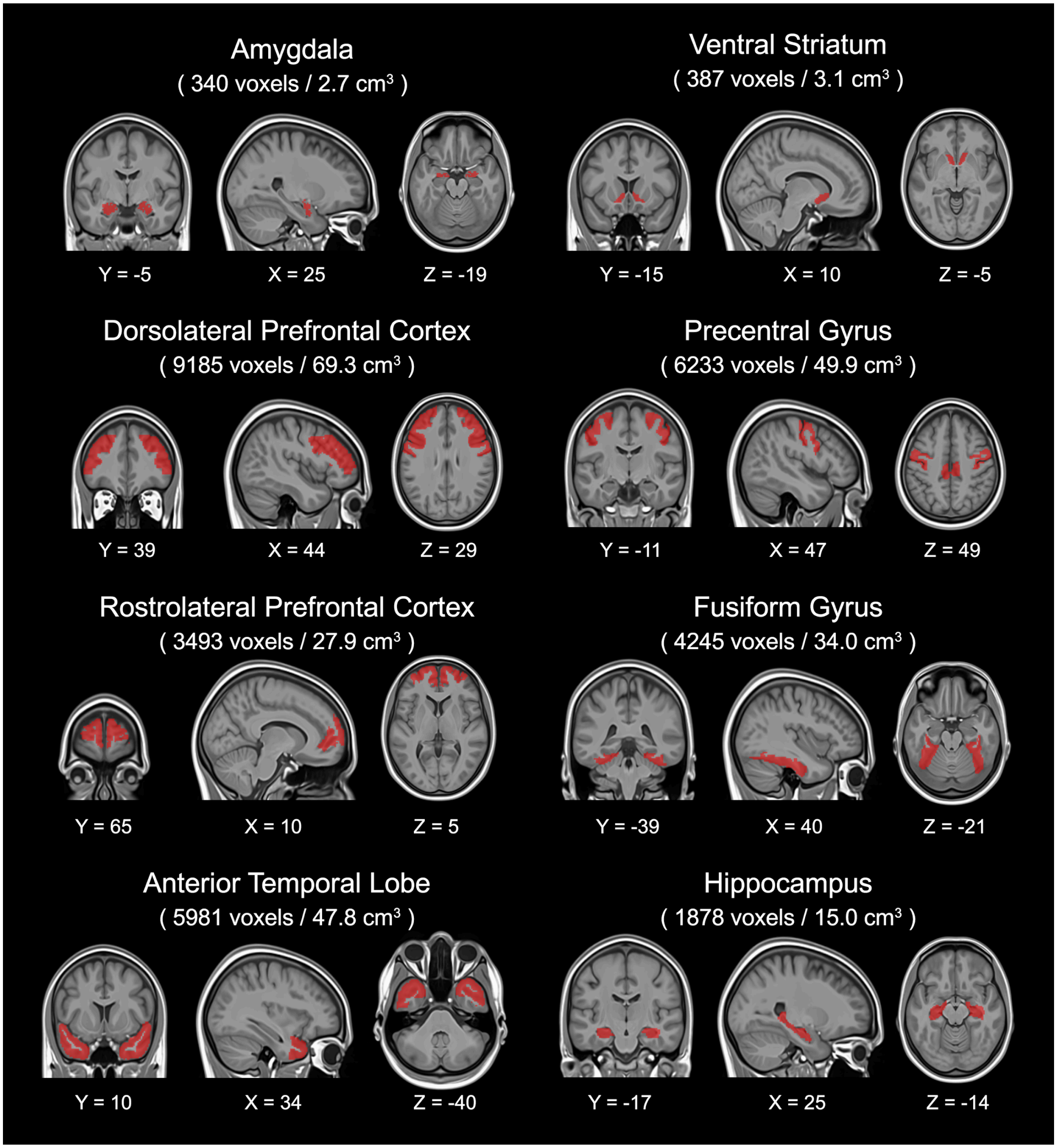

**Fig. S4.** Whole-brain activation maps for the five fMRI tasks not shown in Fig. 4 of the main text.

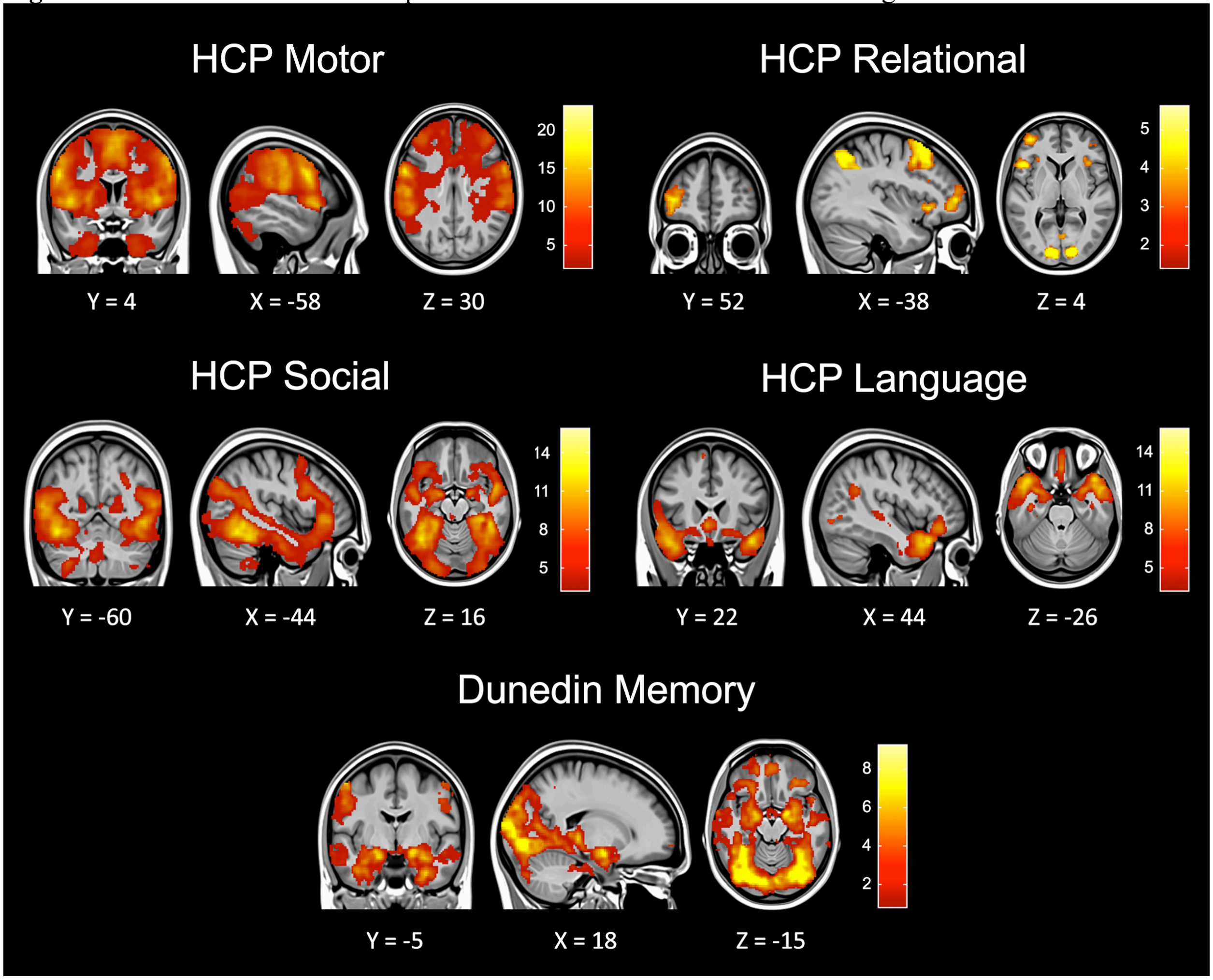

**Fig. S5.** ICCs for response time (RT) and accuracy from fMRI tasks. Each measure is given for the primary experimental condition of interest: the face-matching condition for the emotion tasks, the gain/reward condition for the reward tasks (note that accuracy is undefined for HCP's task), the incongruent/2-back condition for the executive function (EF) tasks, the relation condition for the relational task, the theory of mind condition for the social task, the language condition for the language task, and the encoding condition for the memory task. The fMRI tasks in HCP and the Dunedin Study have widely variable ICCs. Of note, some tasks, like the emotion tasks, were designed to be very easy (i.e. have near ceiling accuracy); this limits between-subjects variance and leads to a low ICC. In this way, the low behavioral ICC was intended.

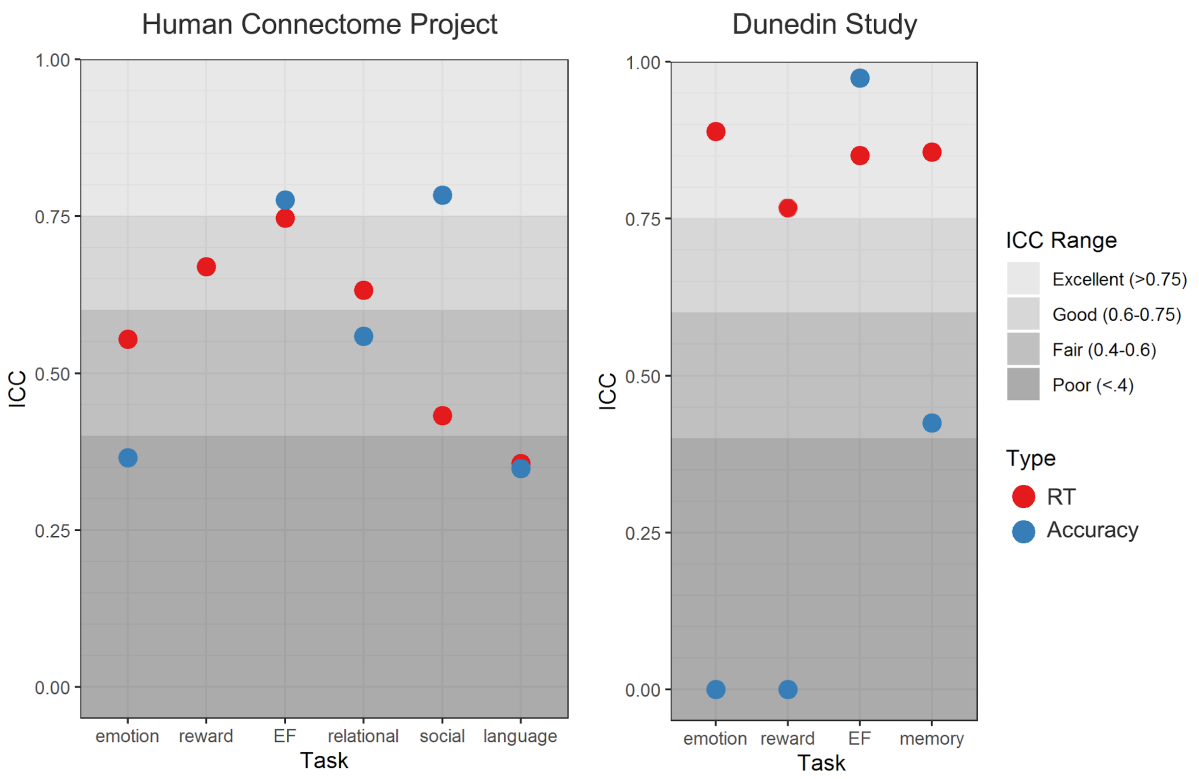

**Fig. S6.** Sensitivity analyses for test-retest reliability of region-wise activation measures in the Human Connectome Project (HCP) and the Dunedin Study. ICCs shown are calculated separately across hemispheres (A and D), within functionally defined ROIs (B and E), and in only the N=26 unrelated individuals in the HCP (C).

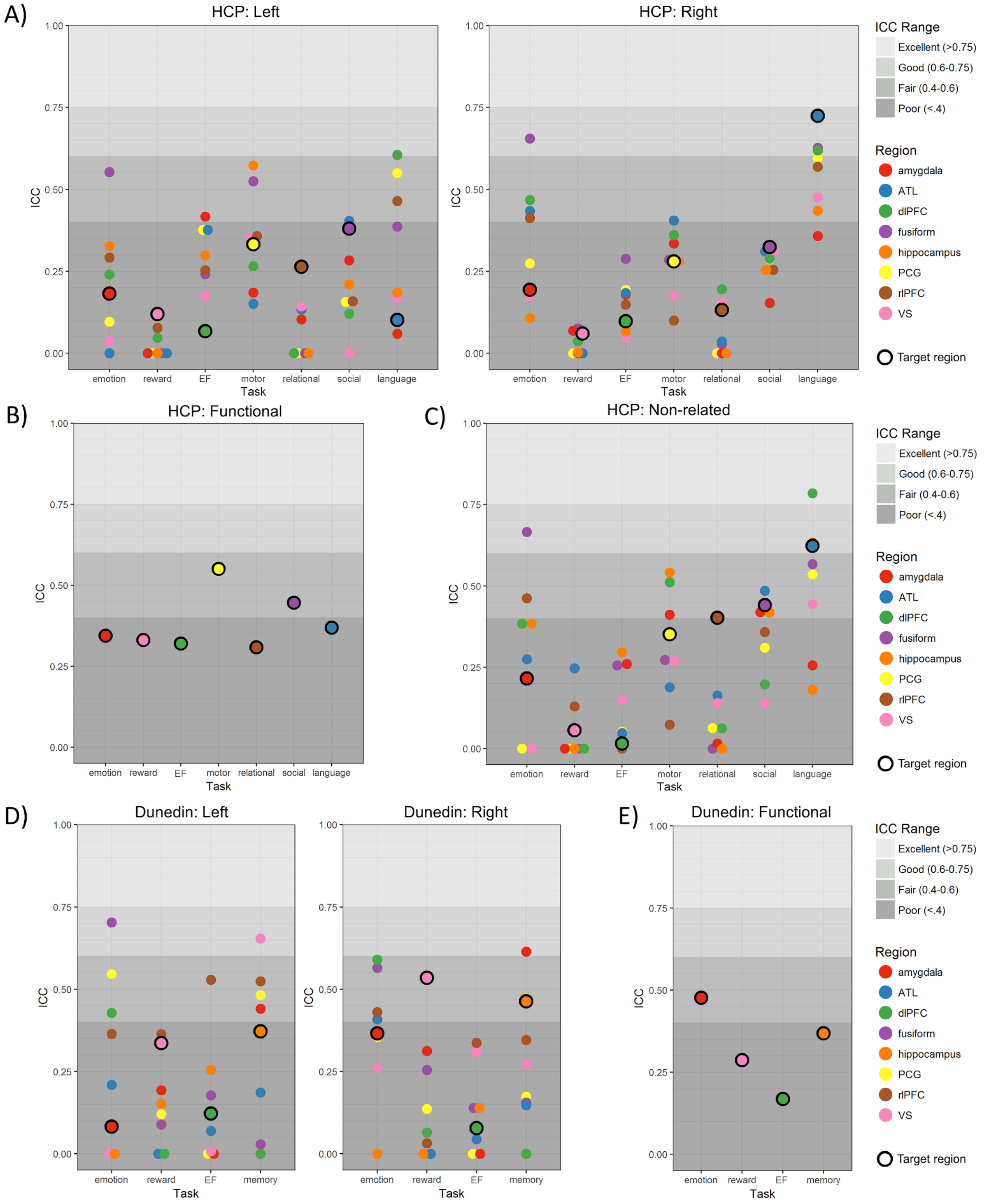

**Table S1.** Moderators tested in meta-analysis

| Moderator | Level | Count | Mean (sd) | Min | Max |
|---|---|---|---|---|---|
| *Number of citations per year* | - | - | 5.03 (4.38) | 0 | 23.7 |
| *Thresholded on ICC* | No | 840 | - | - | - |
| | Yes | 306 | - | - | - |
| *Retest interval (days)* | - | - | 119.8 (242.3) | 1 | 1008 |
| *Sample type* | Healthy | 1036 | - | - | - |
| | Clinical | 110 | - | - | - |
| *Task design* | Blocked | 859 | - | - | - |
| | Event-related | 239 | - | - | - |
| | Mixed | 48 | - | - | - |
| *Task length (minutes)* | - | - | 10.59 (6.45) | .9 | 31 |
| *Task type* | Emotion | 297 | - | - | - |
| | Executive Control | 139 | - | - | - |
| | Language | 12 | - | - | - |
| | Memory | 134 | - | - | - |
| | Motor | 124 | - | - | - |
| | Pain | 177 | - | - | - |
| | Reward | 210 | - | - | - |
| | Sensory | 44 | - | - | - |
| | Social | 9 | - | - | - |
| *ROI type* | Functional | 374 | - | - | - |
| | ICC-based | 152 | - | - | - |
| | Structural | 451 | - | - | - |
| | Structural+Functional | 166 | - | - | - |
| | Whole-brain | 3 | - | - | - |
| | Cortical | 744 | - | - | - |

| *ROI location* | Subcortical | 399 | - | - | - |
|---|---|---|---|---|---|

**Table S2.** Activation in *a priori* anatomical target ROIs. Mean t-statistics are given for all voxels in each bilateral ROI, as well as left and right separately.

*Human Connectome Project*

| **Task (*ROI*)** | **Mean t (bilateral)** | **Mean t (left)** | **Mean t (right)** | **Peak location (x,y,z)** |
|---|---|---|---|---|
| Emotion *(Amygdala)* | 8.897404 | 9.118625 | 8.681328 | 18, -2, -18 |
| Reward *(Ventral Striatum)* | 5.721121 | 5.604154 | 5.831621 | -8, 12, -4 |
| Cognitive Control *(dlPFC)* | 3.264334 | 2.997859 | 3.519485 | 46, 30, 34 |
| Motor *(Precentral Gyrus)* | 7.594384 | 8.417101 | 6.747241 | -58, 4, 30 |
| Relational *(rlPFC)* | -0.23022 | -0.07973 | -0.36461 | -38, 52, 4 |
| Social *(Fusiform)* | 5.356837 | 5.977268 | 4.761796 | -44, -60, -16 |
| Language *(Anterior Temporal Lobe)* | 5.769964 | 6.041165 | 5.49485 | 44, 22, -26 |

*Dunedin Study*

| **Task *(ROI)*** | **Mean t (bilateral)** | **Mean t (left)** | **Mean t (right)** | **Peak location (x,y,z)** |
|---|---|---|---|---|
| Emotion *(Amygdala)* | 6.133734 | 5.877258 | 6.384245 | 18, -5, -15 |
| Reward *(Ventral Striatum)* | 4.20533 | 4.263221 | 4.150639 | 20, -5, -15 |
| Cognitive Control *(dlPFC)* | 3.163896 | 2.440572 | 3.884862 | -48, 23, 21 |
| Episodic Memory *(Hippocampus)* | 2.942571 | 2.794044 | 3.088901 | -12, 13, -5 |

**Table S3.** Test-retest reliabilities of grey matter volumes for 17 subcortical regions

| ***Region*** | ***Human Connectome Project*** | | ***Dunedin Study*** | |
|---|---|---|---|---|
| | ***Left*** | ***Right*** | ***Left*** | ***Right*** |
| Cerebellum | 0.982 | 0.983 | 0.891 | 0.94 |
| Thalamus | 0.889 | 0.875 | 0.919 | 0.971 |
| Caudate | 0.984 | 0.977 | 0.957 | 0.957 |
| Putamen | 0.934 | 0.979 | 0.917 | 0.973 |
| Pallidum | 0.791 | 0.815 | 0.901 | 0.922 |
| Hippocampus | 0.895 | 0.924 | 0.957 | 0.979 |
| Amygdala | 0.917 | 0.914 | 0.918 | 0.931 |
| Accumbens | 0.831 | 0.822 | 0.767 | 0.942 |
| Brainstem | 0.837 | | 0.979 | |
